# Dynamics of blood NAD and glutathione in health, disease, aging and under NAD-booster treatment

**DOI:** 10.1101/2025.02.24.639825

**Authors:** Liliya Euro, Kimmo Haimilahti, Sonja Jansson, Saara Forsström, Jana Buzkova, Anu Suomalainen

## Abstract

Nicotinamide adenine dinucleotide (NAD) and glutathione are vital molecules that control redox-state, enzyme functions and metabolic flux in hundreds of cellular metabolic reactions. High NAD^+^ level occurs in fasting and has been associated with health outcomes in model systems, while low NAD^+^/NADH ratio occurs in specific diseases. Still, the normal, booster-treated or disease-related levels of NADs or glutathiones are not well known in humans. Here, we present a standardized technology for high-throughput quantitative measurement of NAD^+^, NADH, NADP^+^, NADPH, GSH and GSSG from a single whole-blood sample. In healthy population (n=299;18-70 year-olds) redox metabolites follow normal distribution in blood and remain unchanged during aging. NAD-boosting increased 4-6 fold the blood NAD^+^ depending on individual, pointing to need of personalized dose adjustment in treatment trials. In patients with cancer, diabetes or neurodegeneration, NADs and glutathiones showed disease-dependent “redox fingerprints”. The evidence highlights the potential of redox profiling as an indicator of metabolic pathology and as a measure of treatment response.

## Introduction

The physiological redox system and its implications in maintaining metabolic homeostasis in health and degenerative diseases is gaining progressive attention. The key regulators are the nicotinamide adenine dinucleotides (NADs) - the bodily forms of vitamin B3 - and the glutathione forms. Findings of NAD and glutathione changes are reported in genetic metabolic diseases as well as in degenerative, age-related diseases^1,2^. As their levels can be modified by their precursors - vitamin B3 forms for NAD+ and N-acetylcysteine for glutathione - the interest has been high also in the longevity, aging and healthy lifestyle fields. However, knowledge of the normal ranges for all redox metabolites in accessible bodily liquids, such as blood, in healthy human populations has been lacking, largely because of a lack of methodology to assess full redox profiles in a high throughput manner from large population materials.

NAD^+^ forms are the major electron carriers in cellular metabolic reactions. The major NAD(H)-dependent process in living organisms is a multistep conversion of energy of nutrients into the high-energy phosphodiester bond of ATP. Glutathione, one of the most abundant cellular metabolites, maintains cellular environment in reduced state, protecting proteins and iron-sulfur clusters from oxidative stress and is essential for biosynthesis reactions, such as nucleotide synthesis^3^. Both NAD and glutathione forms are used as cofactors and posttranslational modifications in proteins involved in biosynthesis reactions. Cycle of glutathione oxidation and reduction is tightly coupled with reduced phosphorylated form of NAD^+^, e.g. NADP(H)^4^, linking the two redox systems. The redox properties of NADs regulate the direction of hundreds of cellular biosynthetic pathways varying from amino acid, lipid and nucleotide synthesis to folate cycle, detoxification reactions of drugs and reactive oxygen species^5–7^. NAD^+^ has an essential role as a source of ADP-ribose for three main classes of enzymes, each releasing nicotinamide as a byproduct: 1) mono- and poly-ADP-ribosyltransferases (ARTs, formerly PARPs) utilize ADP-ribose to modify proteins post-translationally and get activated during DNA repair processes^8^; 2) glycohydrolases (e.g., CD38, CD73, CD157) hydrolyze NAD^+^ and release free ADP-ribose crucial for cellular Ca2^+^ signaling^9–11^; 3) sirtuins, main sensors for nutrient state, are activated by increased NAD^+^/NADH ratio^12^. NAD^+^ serves also as a substrate for NAD^+^ kinases, which are responsible for synthesizing the phosphorylated form of the metabolite, NADP^+13^. All in all, balance of redox regulators is essential for health across all life forms, and their pools are consumed or imbalanced as a consequence of diseases.

Levels of NADs and glutathiones are dynamic and respond to exogenous and endogenous changes. Vitamin B3 is food-derived and its nutritional deficiency causes a severe disease, pellagra^2^. Pathogenic genetic variants in NAD-synthesis enzymes lead to NAD^+^ deficiency^14^. NAD^+^ deficiency can also be endogenous, a secondary consequence of metabolic disease. For example, mitochondrial muscle diseases cause NAD^+^ deficiency, detectable both in the muscle and blood^15,16^. In these studies, high-dose niacin treatment restored NAD^+^ levels and provided functional benefits in adult patients: remarkable decrease of liver fat, increase of mitochondrial respiration and biogenesis, muscle strength and exercise tolerance^15^.

The possibility to affect NAD^+^ and glutathione levels by treatment using their precursors makes them important to consider in medicine. Past decade was marked by expanding development of NAD-boosters, vitamin B3-forms: nicotinamide, nicotinamide riboside, nicotinamide mononucleotide and reduced forms of these metabolites, NRH^17^ and NMNH^18,19^. They all are derivatives of niacin, the basic form of vitamin B3. Their efficacy in increasing levels of NAD+ is being extensively studied in model systems and humans^15,17,18,20,21^. N-acetylcysteine (NAC) serves as an alternative source for cysteine and glutathione synthesis^3^. While clinical studies using NAD-boosters, glutathione or NAC in different diseases are increasing in number, their implementation as trial end-points and in clinical use is hampered by the lack of knowledge of their ranges in healthy population, especially in the blood, an accessible sample source. Analysis of the redox metabolites from a single blood sample has been a challenge because of the different pH stability requirements of their reduced and oxidized forms, and the high protein concentration of the blood. Reliable quantification of individual NAD metabolites and glutathione can be achieved through liquid-chromatography mass-spectrometry (LC-MS)^22^, cyclic enzymatic assays and high-performance liquid chromatography (HPLC)^23–26^, but these have not been applicable for the scalable analysis of the whole redox set, especially from a single extract.

Here, we report a quantitative measurement and individual relationships of the major cellular redox regulators, NAD^+^, NADH, NADP^+^, NADPH, GSH and GSSG, from the blood of 299 healthy blood donors, upon B3-vitamin supplementation of healthy individuals and in age related diseases. This was enabled by our new method allowing simultaneous analysis of all the six metabolites from a single blood sample.

## Results

### Analysis of NAD and glutathione forms from the whole blood

First, to enable analysis NAD^+^, NADH, NADP^+^, NADPH, GSH and GSSG from large human cohorts, we developed a quantitative measurement of these six metabolites from a single blood sample (Fig. 1a). The extraction from a sample of 100 µl whole blood was done in non-buffered mixture of alcohols heated to induce protein unfolding^27^, releasing non-covalently bound cofactors while preserving redox-sensitive metabolites. Then, each of six redox metabolites is measured in cyclic enzymatic assay system, followed by colorimetric analysis. Glutathione covalently bound to proteins remains in the pellet post-extraction. Denatured proteins are precipitated by cooling and removed via centrifugation, resulting in a clear supernatant. The stability of NAD^+^, NADH, NADP^+^ and NADPH in extraction conditions was verified on solutions of pure compounds using UV-Vis spectroscopy (Fig. 1b,c). To validate extraction protocol of all six metabolites from whole blood we used spiking approach. Spiking experiments involved addition of known amounts of pure NAD and glutathione compounds individually at different stages of the extraction (Supplementary Table 1), followed by quantification using colorimetric assays. These experiments showed that blood plasma has NADH oxidizing activity, while cells have property to degrade added exogenous NADP^+^, NADPH and NAD^+^. In contrast, reduced and oxidized glutathione added to blood samples, cells or plasma remained intact. Analysis of whole blood, its cellular and plasma fractions using developed method showed that NAD metabolites and glutathione reside inside the cells (Fig. 1d-i). Next, we compared performance of our method against liquid-chromatography–mass-spectrometry. Validation was focused on NAD^+^ and NADP^+^ as there is no MS based method quantitatively measuring all six redox metabolites. Comparative experiment showed good concordance for NAD+ and satisfactory for NADP+ (Fig. 1j,k).

**Figure 1.**
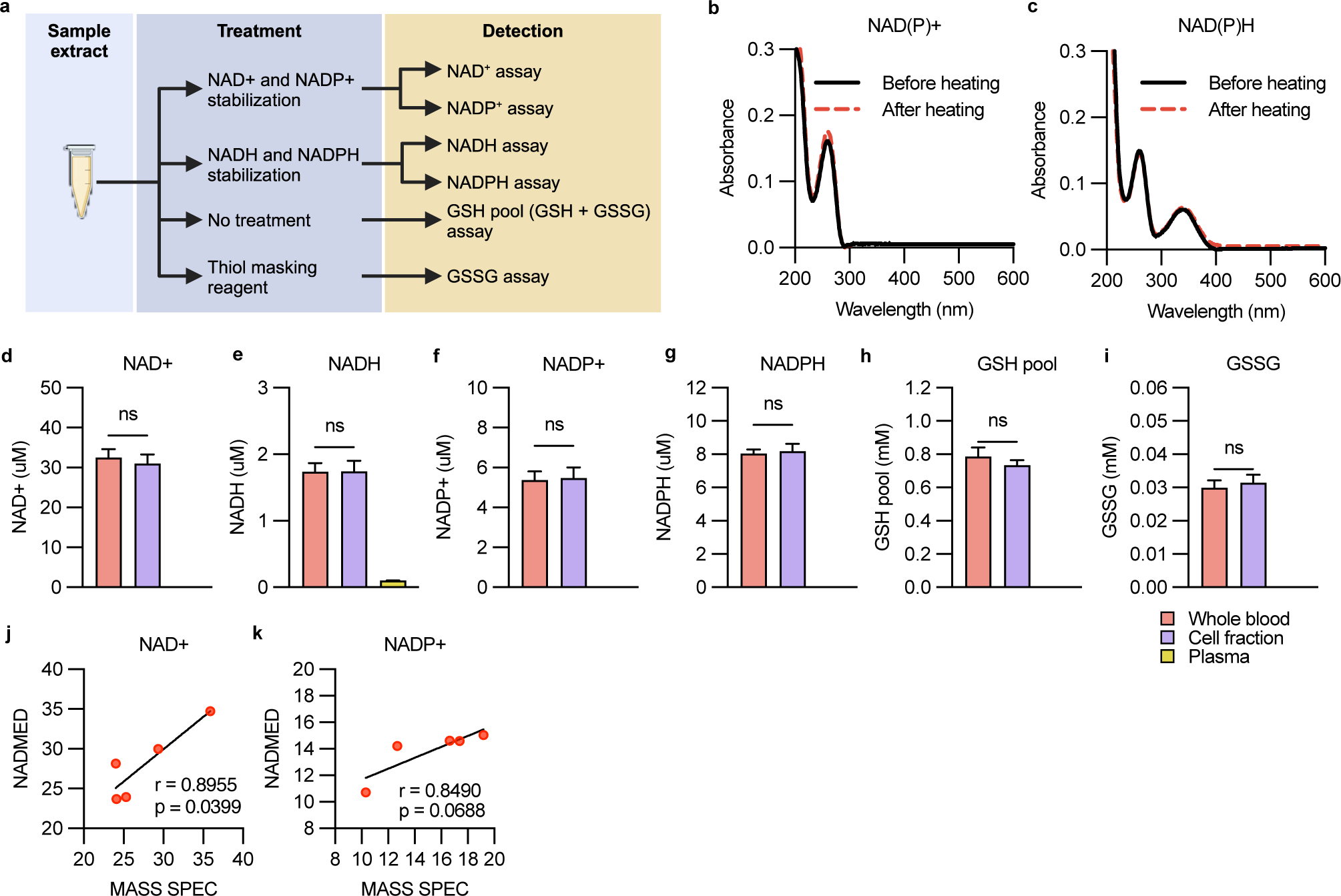
Assay properties. **a.** Scheme for blood sample handling to detect different redox metabolites. **b and c.** Optical spectrum of 10 uM NAD(P)^+^ and NAD(P)H in non-buffered alcohol solution before (black line) and after heating (red dotted line). Light absorption spectra of NAD^+^ and NADP^+^ are identical as are spectra for NADH and NADPH. **d-i.** NAD metabolites and glutathiones in whole blood (n = 2-5), blood cell extract (n = 4-6) and plasma (n = 2-3). Data are mean with standard deviation. **j and k.** Correlation of whole blood NAD^+^ (j) and NADP^+^ (k) between mass spectrometry and NADMED (n = 5, 3 technical replicates per sample). Spearman correlation used for calculating correlation coefficients. Difference between whole blood and blood cell fraction was calculated with Mann-Whitney non-parametric test.

To assess optimal blood handling conditions for reliable measurement of NAD and glutathione metabolites, we investigated their stability in fresh whole blood samples under varying conditions of temperature and time. Over 24 hours at room temperature, NAD^+^ showed tendency to increase by approximately 25% and NADH by 125% from the baseline at collection, while NADP^+^ remained stable and NADPH levels decreased 69% (Supplementary Figure 1a-f). Extending the incubation to 72 hours exacerbated these trends, and even GSH levels fell by 33%, and GSSG increased by 81% from the baseline (Supplementary Figure 1g-l). These findings emphasize the need of swift freezing or analysis of the metabolites, with a maximal time of four hours of exposure to room temperature to prevent significant metabolite dynamics *in vitro*. Next, we investigated effect of freeze-thaw cycle on blood NADs and glutathione concentration. While freezing did not impact the concentrations of NAD^+^, NADH, GSH, or GSSG, it triggered the conversion of NADPH to NADP^+^, with about 50±8% of NADP^+^ originating from NADPH oxidation (Supplementary Figure 1m-r). To minimize such unwanted events, we developed a standardized thawing procedure for frozen samples, with a small sample volume (150-200 µL): thawing under strict cooling conditions (+0°C to +2°C) within 12-15 minutes in an ice-water bath, without agitation. Following this protocol, the developed redox profiling method demonstrated high precision with intra-assay coefficients of variation (CV%) for each redox metabolite measured from frozen blood samples (Supplementary Tables 2 and 3). The sensitivity of the quantification is high: 0.35 µM for NAD^+^ and NADP^+^, 0.28 µM for NADH and NADPH, 72 µM for the GSH pool and 10 µM for GSSG (Supplementary Figure 1s). These limits are well below the physiological concentrations of these metabolites in whole blood (Fig. 1 and 2), supporting the utility of our methodology for quantitative redox profiling in human whole blood samples.

**Figure 2.**
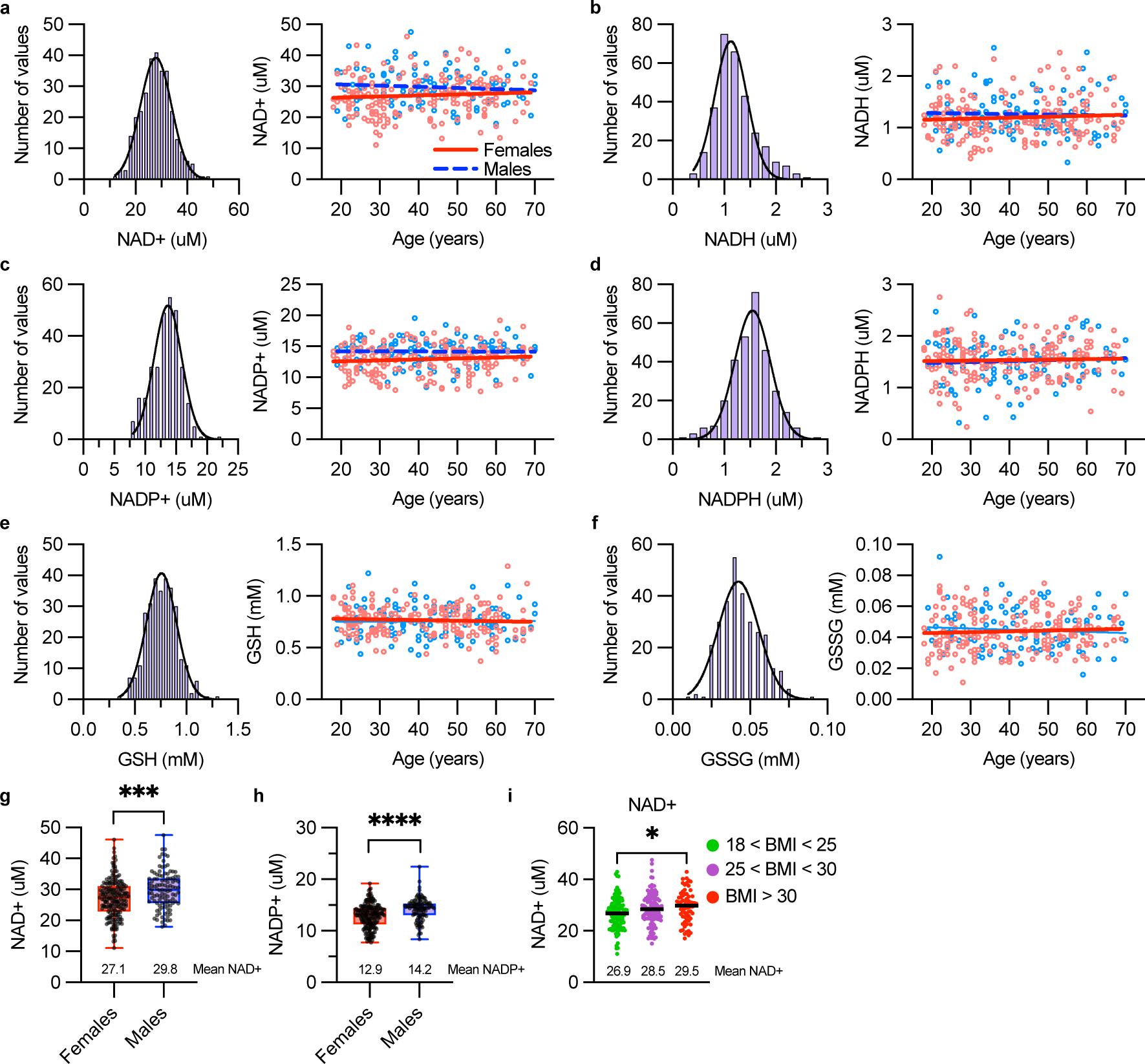
Blood levels of NAD metabolites and glutathiones measured in healthy population remain stable during aging and follow normal distribution. **a-f.** Frequency distribution and levels of NAD+ (a), NADH (b), NADP+ (c), NADPH (d), GSH (e) and GSSG fF) in the whole blood of 299 healthy donors at the age of 18-70 years. During extraction of GSH and GSSG one sample was lost. Red and blue lines represent linear regression fit of the data separately for females and males. Black line represents Gaussian distribution. **g and h.** Blood NAD+ and NADP+ in females and males. **i.** Effect of BMI on blood NAD+ level. BMI was not available from two individuals and they were excluded from the analysis. Gender differences were analyzed with unpaired parametric t test. For BMI effect one-way ANOVA with Dunnet’s multiple comparison test was applied. *p < 0.05, ***p < 0.001, ****p < 0.0001

### Establishment of normal concentrations of NAD and Glutathione forms in Whole Blood

Next, we applied our profiling methodology to define human population values of the six redox metabolites using frozen whole blood samples from 299 anonymous healthy donors (191 females and 108 males, aged 18-70 years) provided by the Finnish Blood Service BioBank and 38 healthy in-house volunteers. Sample handling and storage details are summarized in methods section. We observed that biobank samples (frozen within 3-17 hours) and fast-frozen in-house samples (within ∼1 hour) showed slight variations in metabolite levels. Specifically, in-house donors exhibited lower NAD^+^, NADH and NADP^+^ levels (on average 3.2, 0.47 and 1.6 uM lower, respectively) but higher NADPH levels (on average 0.88 uM) compared to biobank samples (Supplementary Figure 2a). Glutathione levels remained consistent between the two cohorts, in agreement with previous observations about storage conditions (Supplementary Figure 1). To account for storage time effects, we combined the Red Cross and in-house samples to obtain normal value ranges for frozen blood samples (n=337). These ranges were derived from mean ± standard deviation (Table 1). The control values are also valid for NAD^+^, NADH, GSH and GSSG in fresh blood (within 72 hours of storage at +4°C). However, NADP^+^ and NADPH levels in frozen blood differ from fresh blood due to NADPH oxidation during the freeze-thaw cycle. Fresh blood remains the most suitable sample type for analyzing physiological levels of phosphorylated metabolites. Notably, 50% of measured NADP^+^ in frozen blood originated from NADPH, allowing for extrapolation of NADPH and NADP^+^ levels before freezing, if fresh sample analysis is not feasible. We estimated physiological NADP^+^ and NADPH levels in our healthy cohorts, based on frozen sample results, and validated the theoretical calculation by analyzing fresh whole blood samples from 12 additional healthy volunteers (Supplementary Table 5).

**Table 1.**
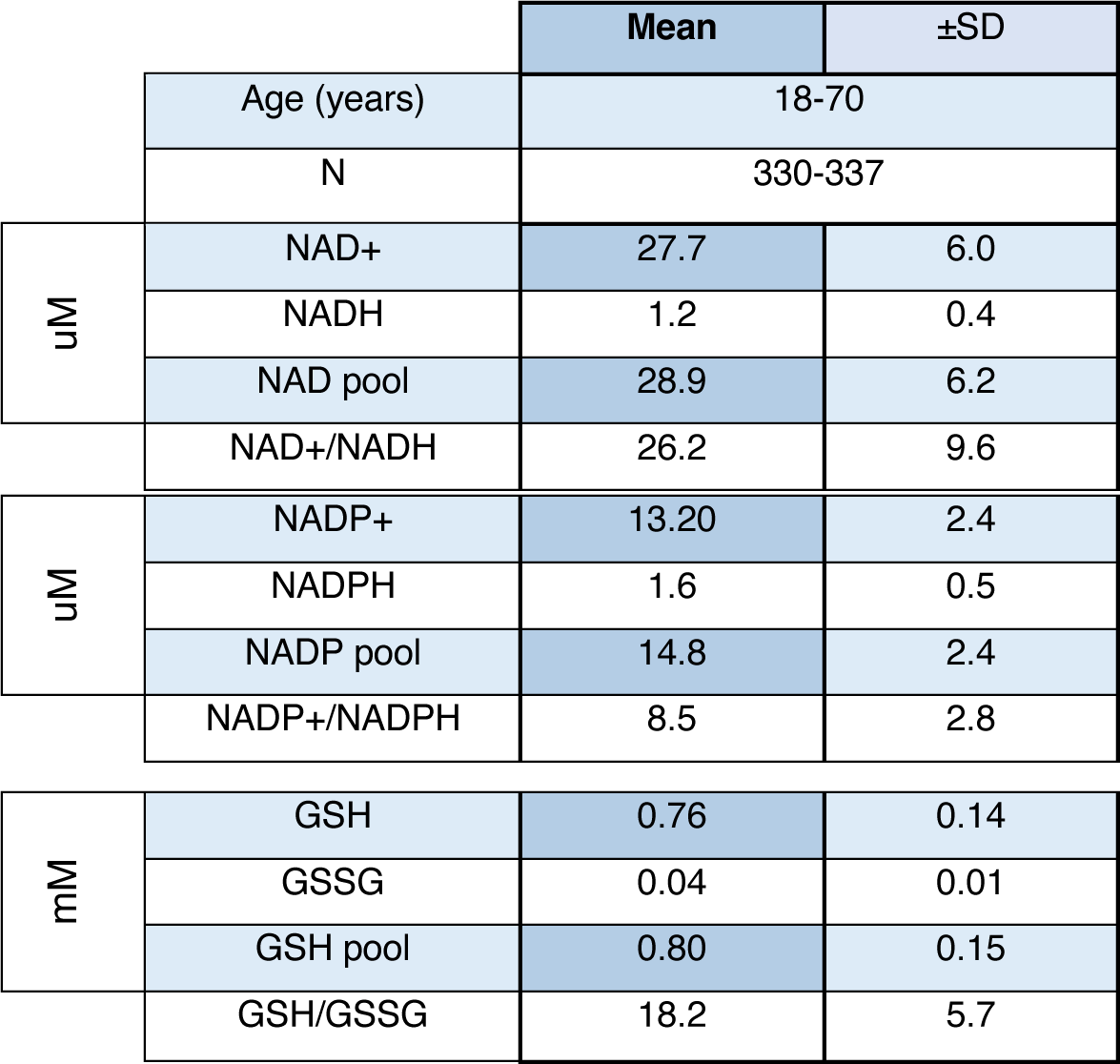
Control ranges for all measured metabolites from frozen blood samples of 299 RedCross blood donors and 38 in-house controls. For GSH and GSSG 298 samples from Red Cross donors and 32 samples from in-house controls could be analyzed. Data are mean ± standard deviation.

### Redox-metabolite concentrations follow normal distribution

The metabolite distributions in the blood samples typically followed a normal distribution across both genders, with no significant age-related changes in the whole population (Fig. 2a-f; Supplementary Figure 2b-g). When grouped by age and gender, female subjects at the age of 31-40 years had slightly higher NAD pool and GSH/GSSG ratios compared to 18-30 and 41-50 year-old females, respectively (Supplementary Figure 2j,s). In the whole population, NAD^+^ and NADP^+^ were higher in males than females (on average 2.7 and 1.3 µM, respectively; Fig. 2g,h). In the subgroup analysis, this difference was clear in young and middle-aged individuals (Supplementary Figure 2h,l). Young males displayed also higher NADP^+^/NADPH ratio in frozen blood compared to age-matched females, while NADH and GSSG were higher and GSH/GSSG ratio lower in 31-40 years old males compared to females (Supplementary figure 2i-s). BMI did not differ between males and females, thus not explaining the observed differences between genders (Supplementary figure 2t). BMI >30 associated with an average increase of 2.6 µM in NAD^+^ levels in both genders (Fig. 2i). Overall, no major changes of mean concentrations of blood NAD or glutathione metabolites occurred during aging, but subgroups who had lowered levels existed in all ages.

Together, the results show that our sensitive assay enabled establishment of NAD and glutathione values in a large healthy population cohort, serving as a basis to interpret normal and abnormal values in whole-blood samples.

### 500-750 mg niacin per day saturates steady-state blood NAD^+^ in healthy subjects

Next, we explored the impact of NAD^+^ precursor niacin on blood NAD and glutathione metabolites in eight individuals, who were healthy controls in our previous clinical pilot study using niacin as NAD^+^ booster for mitochondrial myopathy patients that showed lowered NAD^+^ values^15^. They ingested increasing doses of niacin (250-1000 mg daily) for 16 weeks as described previously^15^, and the blood samples were taken in different timepoints during the supplementation. Niacin resulted in considerable dose-dependent increases in erythrocyte NAD^+^ concentrations, with also subject-dependent dynamics of NAD^+^ increase. At 16 weeks of treatment with 1000 mg dose the levels of NAD^+^ increased four to six-fold (Fig. 3a) and NADH levels elevated as well but showed large inter-individual differences (Fig. 3b). NADP(H) and glutathione remained largely unchanged, with NADPH and GSH showing transient increase after 500-750 mg of niacin (Fig. 3c-f). Figure 3g summarizes the maximal saturation levels of all six metabolites. Erythrocyte NAD(H) levels started to saturate already with 500-750 mg of niacin and increasing the dose up to 1000 mg had only a slight additional effect on steady-state NAD^+^ in blood cells. These results show that to reach a specific steady-state level of blood NAD^+^, different individuals required different dosages.

**Figure 3.**
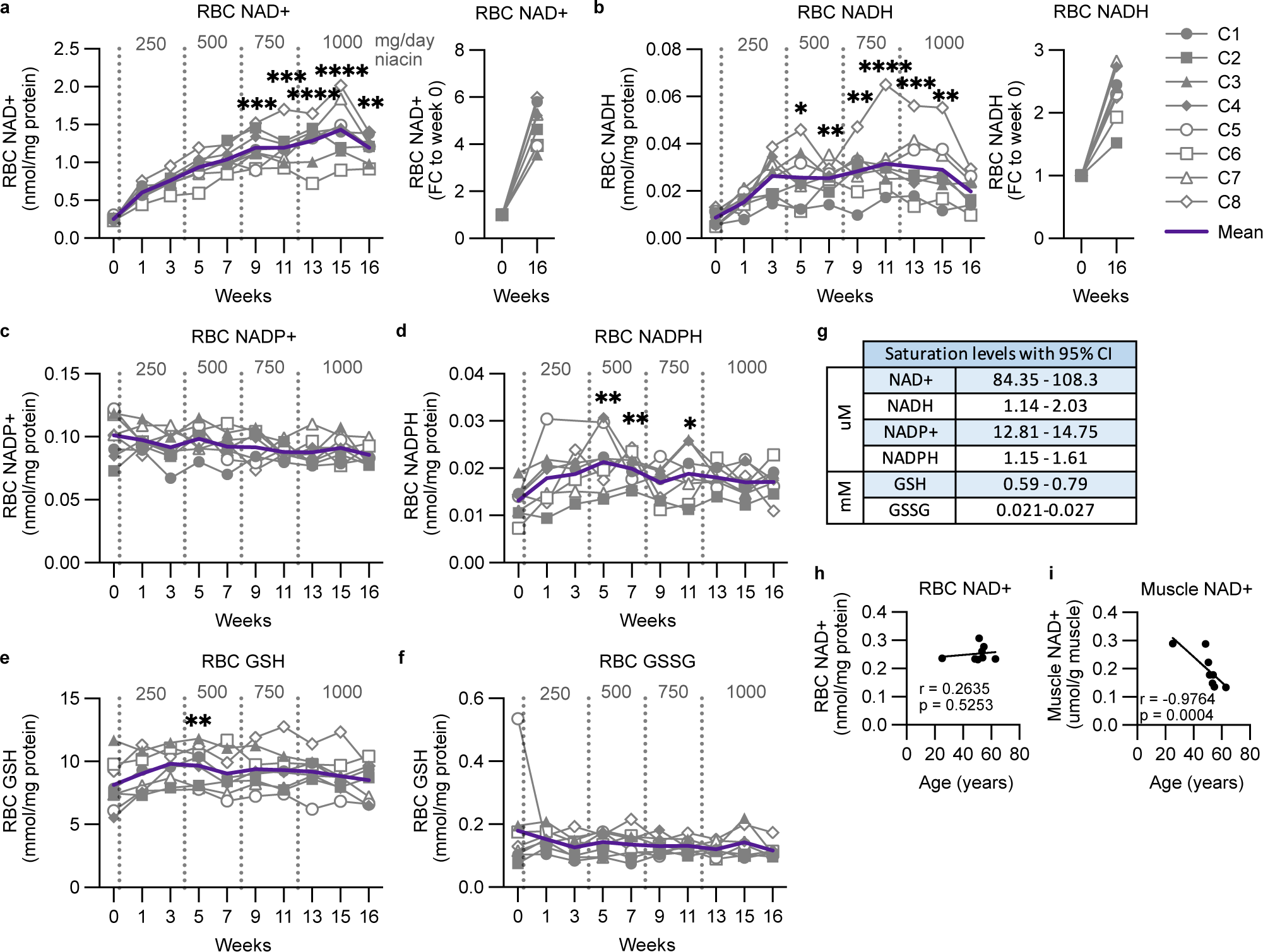
Red blood cell NAD metabolites and glutathione saturate upon supplementation with niacin in healthy individuals. **a-f.** Red blood cell NAD+ (a), NADH (b), NADP+ (c), NADPH (d), GSH (e) and GSSG (f) levels of eight healthy individuals during niacin supplementation. **g.** Saturation levels of each NAD metabolite and glutahtione during niacin supplementation. Values calculated for whole blood (see Methods). **h and i.** Spearman non-parametric correlations of age with red blood cell (h) and muscle (i) NAD+. Muscle NAD+ was analyzed with mass spectrometry. Statistical analysis in a-f was performed with Friedman non-parametric ANOVA with Dunn’s multiple comparison test, p values against week 0. Data from week 3 was not included into the statistical analysis due to a missing data point. *p < 0.05, **p < 0.01, ***p < 0.001, ****p < 0.0001

Similar to what we found in the large cohort of the whole blood analyses (Fig. 2), NAD^+^ in erythrocytes of the eight study subjects did not correlate with age (Fig. 3h). However, their muscle NAD^+^ levels tended to decrease with age (Fig. 3i).

### Specific NAD and glutathione forms correlate with glucose and lipid homeostasis and liver function

The result that the redox metabolites follow a gaussian distribution in normal population, together with the previous knowledge that niacin supplementation can increase blood glucose and decrease cholesterol levels^28^ prompted us to ask whether physiological NADs and glutathione levels correlated with glucose and lipid homeostasis, liver function tests, B vitamins and fatty acid metabolism in healthy individuals (Fig. 4). NAD^+^ correlated positively with glycosylated hemoglobin (HbA1c), a measure of long-term increased blood glucose homeostasis (Fig. 4a). During niacin treatment, fasting glucose, insulin and HOMA1-IR (insulin resistance score based on blood glucose and insulin) increased in some individuals up to 1.9-fold after niacin dose of 500 mg, suggesting increased insulin resistance (Fig. 4b-d). Of metabolic biomarkers, metabokine GDF15 correlated positively with NAD^+^, while FGF21 showed positive correlation with NADH (Fig. 4a). At baseline, HDL cholesterol metrics (total HDL and HDL3) and apolipoprotein A1 (ApoA1) showed positive correlation with NADPH (Fig. 4a). Liver is the major site of HDL synthesis and interestingly NADP^+^ correlated positively with liver markers alanine transaminase (ALT), aspartate transferase (AST) and gamma-glutamyl cysteine (Fig. 4a). Erythrocyte glutathione forms showed less correlation with circulating factors, but GSSG correlated with NADP^+^ and liver markers including gamma-glutamyl cysteine, also a glutathione precursor (Fig. 4a).

**Figure 4.**
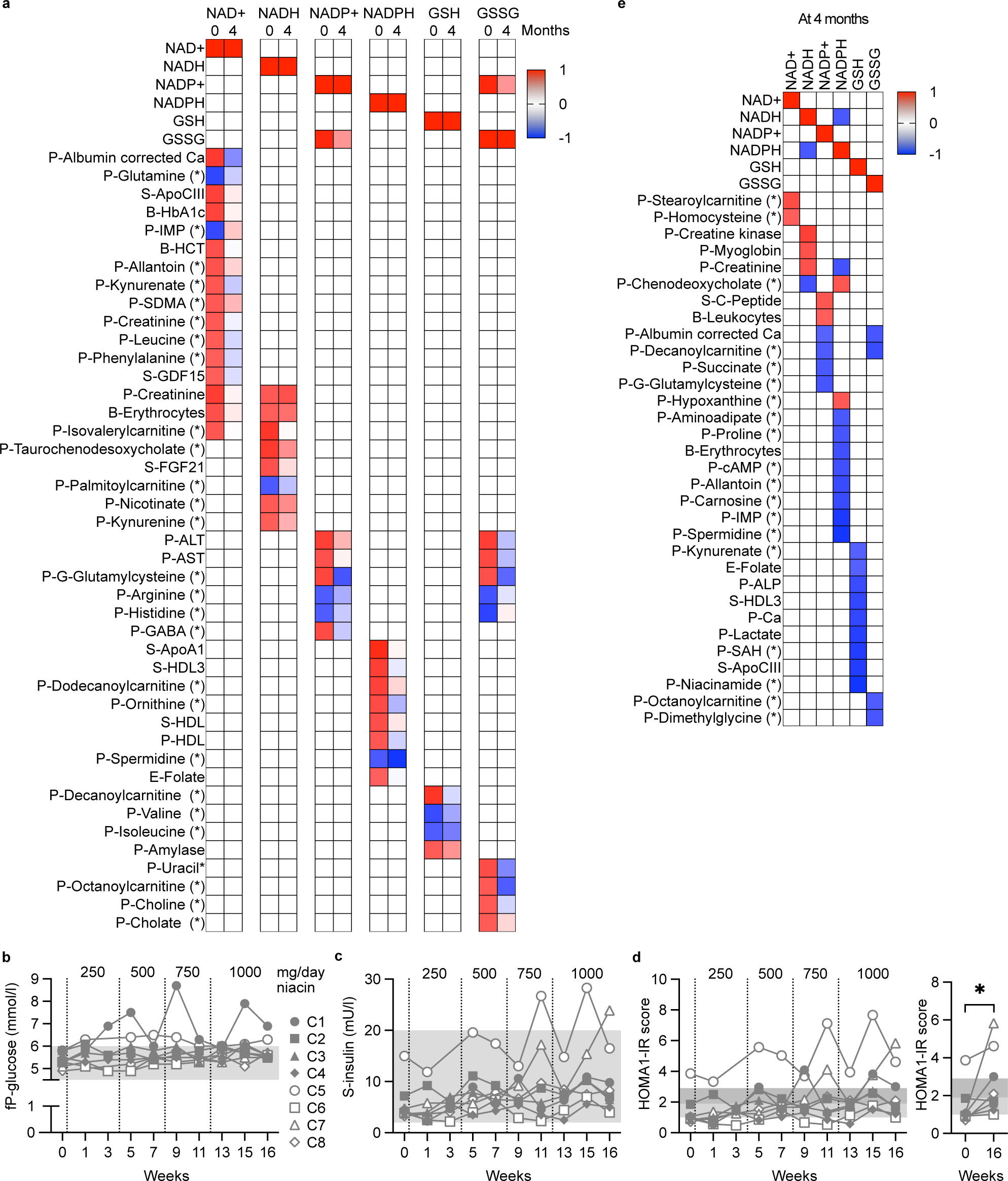
Glucose and lipid parameters correlate with NAD metabolites. Data are from the eight individuals who received niacin with increasing doses for 4 months. **a.** All significant correlations of RBC NAD metabolites and glutathiones with blood metabolites at baseline and their correlations after 4 months on niacin. **b-d.** Response of fasting blood glucose (b), insulin (c) and HOMA1-IR (d) to different doses of niacin. **e.** Correlations of NAD and glutathione metabolites after 4 months on niacin treatment. Spearman non-parametric correlation analysis used as statistical test. All reported correlations in panels a and e have r value over 0.4 and p value under 0.05. In panels b-d, light gray area indicates the normal reference intervals and in the panel d the dark gray area indicates early insulin resistance (scores 1.9-2.9). Wilcoxon matched-pairs signed rank test used in panel d. Sample type: P, plasma; S/not indicated, serum; B, whole blood; RBC, red blood cells. Ca, calcium; ApoA1/CIII, apoliprotein A1/CIII; HbA1c, glycosylated hemoglobin; IMP, inosine monophosphate; HCT, hematocrit; SDMA, symmetric dimethylarginine; GDF15, growth differentiation factor 15; FGF21, fibroblast growth factor 21; ALT, alanine aminotransferase; AST, aspartate aminotransferase; GABA, gamma-aminobutyric acid; cAMP, cyclic adenosine monophosphate; ALP, alkaline phosphatase; SAH, S-adenosylhomocysteine.

Most of the observed correlations at baseline disappeared after 4 months on niacin treatment (Fig. 4a). However, NADH remained positively correlated with creatinine and positive correlations also with other muscle metabolism markers creatine kinase and myoglobin appeared (Fig. 4a,e). Different to baseline, at 4 months NADPH associated with several metabolite of purine metabolism, such as hypoxanthine, inosine monophosphate (IMP), cyclic adenosine monophosphate (cAMP) and allantoin (Fig. 4e).

The study demonstrates the relationship of specific redox metabolites to common clinical parameters.

### Blood NADs and Glutathione Metabolites are Perturbed in Various Diseases

To explore whether the redox pools were modified by chronic age-related diseases, we examined blood NAD and glutathione metabolites across 11 different diseases, including various cancer types, diabetes (Type II) and neurodegenerative diseases (Parkinson’s and Alzheimer’s). A total of 164 frozen blood samples were sourced from the Auria BioBank in Turku, Finland, with the majority collected provisionally at the time of establishing a diagnosis or treatment initiation. However, we did not have information of the disease severity at the sampling time (Supplementary Table 6, Fig. 5 and 6). The heatmap summarizes the association of different metabolites with the diseases (Fig. 5a).

**Figure 5.**
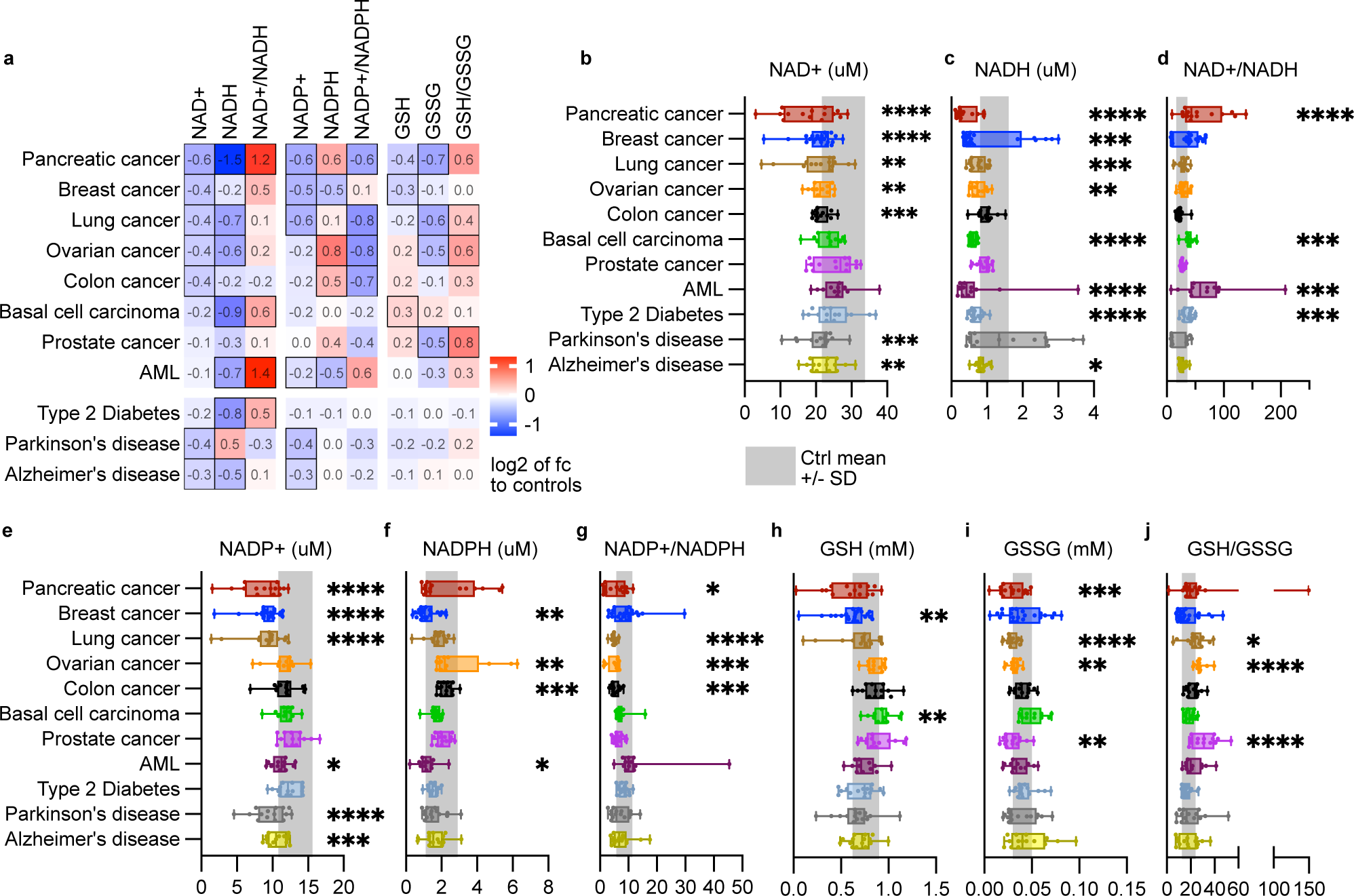
Whole blood levels of NAD metabolites, glutathione and their ratios in different diseases. **a.** Whole blood metabolite and ratios compared to control population (n = 330-337). Data mean fold change compared to controls and adjusted for log2. All statistically significant differences outlined with black borders. **b-j.** Absolute whole blood metabolite levels and ratios in different disease types. Gray area represents control range (mean ± SD) defined in Table 1. Statistical analysis with Kruskal-Wallis non-parametric ANOVA with Dunn’s multiple comparison test, comparison against control population. *p < 0.05; **p < 0.01; ***p < 0.001; ****p < 0.0001. AML, acute myeloid leukemia.

**Figure 6.**
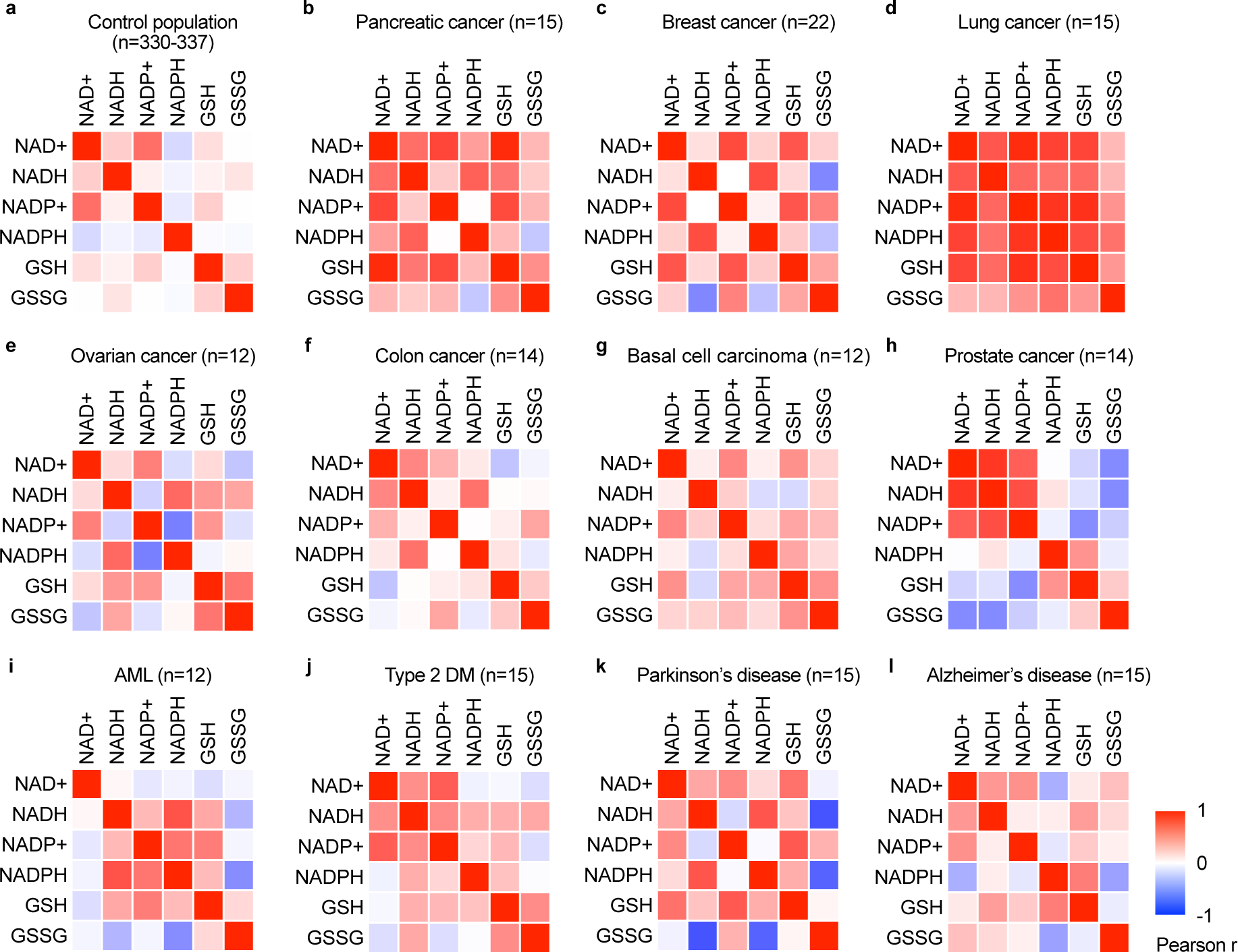
Redox fingerprints are distinct to control population and differ between the diseases. **a-l.** Redox fingerprints, thus heatmaps of Pearson correlation coefficients, of NAD(P)(H) and glutathione profiles in the indicated patient groups. AML, acute myeloid leukemia.

The NAD^+^ and NADH values were low or within the lower end of the healthy range especially in the pancreatic, breast, lung and ovarian cancer (Fig. 5b,c). As NAD^+^ is the precursor of NADH, the low levels of both metabolites suggest NAD^+^ depletion while the NAD^+^/NADH ratio remained unchanged in most of the diseases (Fig. 5d). While suggestive subgroups were visible in e.g. breast cancer, the number of samples and lack of disease-stage data hindered subgroup division. Parkinson’s and Alzheimer’s diseases showed both low NAD^+^. Of note, blood NAD metabolites were changed in all studied diseases, except for prostate cancer, while glutathione metabolites demonstrated alteration only in cancers (Fig. 5b-j). In line with NAD(H) levels, NADP^+^ levels were decreased in several conditions including pancreatic, breast and lung cancers, acute myeloid leukemia (AML) and Parkinson’s and Alzheimer’s disease (Fig. 5e). Unlike other NAD metabolites, blood NADPH levels were increased in most of the cases except for breast cancer and AML (Fig. 5g).

### Correlation heatmaps reveal pathology-related redox fingerprint patterns

Prompted by the redox metabolite changes in different disease groups, we asked whether the ratios between the metabolites differed from healthy controls. The correlation heatmaps – redox fingerprints – (Fig. 6a-l) visualized the changes in redox state compared to the controls. In most of the diseases NAD^+^ was strongly positively associated with NADH and NADP+, which is in line with known role of NAD^+^ as substrate for NADH and NADP+ synthesis. As an exception, in AML which is a blood cell cancer, NAD^+^ did not correlate with any other redox metabolites, potentially because of high number of white blood cells (Fig. 6i). While NAD metabolites did not correlate with glutathione in controls (Fig. 6a), in most of the diseases a correlation was observed. The fingerprints are not disease specific, but healthy and diseased fingerprints are remarkably different.

## Discussion

Here, we report a standardized method for analyzing redox profiles in the whole blood, enabling measurements of all the six redox metabolites (NAD^+^, NADH, NADP^+^, NADPH, GSH, GSSG) from fresh or frozen blood, from a single sample. Our findings demonstrate that whole-blood NAD and glutathione levels follow normal distribution in healthy individuals and remain unchanged during aging. Analysis of biobank samples from patients with different diseases indicates that a wide range of diseases modify remarkably the redox metabolite levels and their ratios. Especially a set of solid cancers and neurodegenerative disease patients (Parkinson’s) showed lowered NAD-metabolite values and in some cancers also glutathiones were changed. Our finding that several diseases show lowered blood NAD^+^ levels and overall imbalanced redox metabolites is intriguing, as it suggests that a variety of disease processes increase NAD^+^ usage, potentially by activated repair processes including PARP enzymes a major user of NAD^+8^.The results have high potential relevance for clinical and research applications, follow-up of metabolic interventions and detection of disease-induced NAD^+^ or glutathione deficiency to select patients for treatment.

Our study was motivated by the previous observation that mitochondrial muscle diseases cause a secondary NAD^+^ deficiency in the skeletal muscle^15^. Importantly, NAD^+^ was systemically lowered, detectable in the blood of the patients. Restoration of the NAD^+^ levels by niacin, a NAD^+^ precursor and B3 vitamin, provided health benefits, slowly reversing physical performance of some adult patients^15^.To find patients who have NAD^+^ deficiency and could benefit from B3 treatment, we needed to set up validated technology and assess their values in the normal population. The method described here measures NAD^+^, NADH, NADP^+^, NADPH and reduced and oxidized glutathione quantitatively and accurately from a single blood sample, using a non-destructive extraction of target metabolites. Liquid-chromatography mass spectrometry (LC-MS) is considered to be a gold standard for redox metabolite measurements^41^. While it is accurate, it is also costly and relatively slow, limiting its use for large sample materials. Also, the reduced NADH and NADPH forms are volatile in the LC-MS setting due to composition of the mobile phase, making their detection by LC-MS complicated. Furthermore, varying pH stability requirements of the redox metabolites^37,38^ and the nature of whole blood as a sample, rich in proteins and heme, add to the complexity of the analyses. Here we show that the integrity of the redox metabolites can be preserved and that the NAD and glutathione pools reside primarily in the cellular compartment of the blood sample, with only trace amounts in the plasma. Our separate assessment of each metabolite using optimized enzymatic assays was calibrated against validated standards and LC-MS and allows high-throughput analyses. The ability to measure oxidized and reduced forms of NADs and glutathiones from the same sample also minimizes effects of separate sample handling. This technology allowed us to perform comprehensive investigation of properties of target metabolites in the blood as well as to assess their levels in different population cohorts.

We show that healthy subjects have stable NAD and glutathione concentrations and their ratios across different age groups, from 18 to 70 years. In accordance with a previous study, males had higher blood NAD^+^ than females^29^, and the same was true for NADP^+^. Our small cross-sectional analysis of parallel blood and muscle samples from a set of healthy subjects showed that muscle NAD^+^ decreases during aging, while blood levels were stable. These results are in line with a previous report from the human muscle^48^. In mitochondrial muscle disease, however, both muscle and blood showed decreased NAD^+^ levels^15,30^ suggesting that blood is a valid source of redox-metabolite analysis in the context of pathology. Considering clinical utility, subtle decline in aging tissues but not in the blood in normal subjects suggests that a decline in systemic blood circulation reflects a chronic disease setting.

We found that niacin intake can elevate blood NAD^+^ levels to a maximum of 80-120 µM, a four to six-fold increase from normal levels, reaching a plateau in blood with daily doses of 500-750 mg. The other NAD forms showed only modest increases in response to niacin, suggesting differential regulation of individual metabolite pools in steady state. Chronic use of NAD^+^ boosters is not without risks, as previously shown for cardiovascular events in in a large retrospective epidemiological study^15,30^. While niacin supplementation can improve muscle mass and positively impact lipid profiles^15^ in mitochondrial disease patients and healthy population, it has been also associated with hyperglycemia^28^. In our study, blood glucose and insulin resistance started to increase with niacin doses over 500 mg/day. Even in healthy controls without any supplementation blood glucose and insulin resistance correlate positively with NAD metabolites. Our findings indicate that glucose metabolism should be followed up if long-term NAD-interventions are needed. Also, in patients with elevated NAD^+^ (such as those suffering from metabolic syndrome^31^), NAD^+^ booster use needs careful consideration due to potential exacerbation of insulin resistance.

Modified NAD and glutathione metabolism is associated with a number of diseases, but no data exist of how specific pathology is shifting blood redox profile. Our comprehensive redox profiling across various cancer types and age-related neurodegenerative diseases revealed modified “redox fingerprints”, major changes in the correlative patterns of the metabolites with each other, differing from the healthy controls. The findings stimulate further studies in large biobank samples to clarify whether such signatures could prove valuable for detecting pathology or follow-up of treatment effect in clinical trials.

In conclusion, our study provides a robust methodology for redox profiling and establishes baseline data for NAD and glutathione metabolites in healthy populations. The observed dynamics of these metabolites in response to supplementation and disease states offer new insights into redox biology and metabolism. These findings have significant implications for the development and monitoring of NAD-targeted therapies, emphasizing the need for personalized approaches in metabolic interventions.

### Limitations of the study

The normal levels were established using Finish RedCross samples from assumingly healthy volunteers upon blood donation, and clinical pathology samples were collected from Auria Biobank material. Both biobanks followed similar sample processing protocols, aligning with standard clinical laboratory practices for the time between sampling and cooling/freezing. Normal levels for NAD^+^, NADH, GSH pool and GSSG were determined from frozen blood samples of healthy donors. These values are considered valid for both frozen and fresh blood samples. However, the levels of NADP^+^ and NADPH in fresh and frozen blood differ. We estimated NADP^+^ and NADPH levels in fresh blood based on concentrations measured in frozen samples. It is based on assumption that 50% ± 8% of the NADP^+^ content in frozen blood accounts for NADPH that oxidized during the freeze-thaw cycle.

## Author contribution

LE - developed the method, designed the study, performed measurements, analyzed the data, wrote the article; KH – analyzed the data, prepared the figures, wrote the article; SJ – performed measurements, contributed to article writing; SF – contributed to article writing, JB – designed the study, organized sample acquirement from BioBanks, contributed to article writing; AW – designed the study, analyzed the data, wrote the article.

## Acknowledgements

We express our special gratitude to Markus Innilä, Satu Malinen, Liina Lassila and Babette Hollmann for their technical assistance. We thank Dr. Eija Pirinen for constructive discussions on interpretation of redox profiling results for retrospective samples from published clinical trial. FIMM Metabolomics facility is acknowledged for cross-check experimental validation of NADMED method measuring NAD^+^ and NADP^+^ concentrations in blood. The Blood Service Biobank personnel are acknowledged for organizing research project-specific sample collection in blood donation activities and Aurea biobank thanked for samples of diseased individuals. Open resource Biorender.com is acknowledged for enabling creation of Fig. 1a and part of Supplementary Table 1.

## Ethical statement and Biobank agreements

Ethics committee of the hospital district of Uusimaa and Helsinki gave ethical approval for collection of the in-house control blood samples. In-house volunteers who donated blood for test development and validation were from our local community and recruited through an advertisement. All participants gave written consent prior blood donation and samples were anonymized. The human samples were collected by Red Cross blood donor service and Aurea biobank according to ethical permits of the biobanks and shared based on research collaboration to this study.

## Materials and Methods

### Principle of developed redox profiling method

For quantitative analysis of NAD^+^, NADH, NADP^+^, NADPH, GSSG and GSH in a single blood sample, we developed a comprehensive method, “NADMED”, consisting of novel extraction coupled with detection based on previously published cyclic enzymatic reactions with colorimetric detection^23,24,32–34^ in a high-throughput 96-well-plate analysis. Commercially available pure compounds of NADs and glutathiones were used for preparation assay standards. Concentration of pure compounds in Standard stocks were validated by UV-VIS spectroscopy as described elsewhere. Briefly, the extraction uses non-buffered alcohol solution mixture, heated to induce protein unfolding and releasing cofactors while preserving redox-sensitive metabolites. Denatured proteins are precipitated by cooling and removed via centrifugation, keeping of NAD^+^, NADH, NADP^+^, NADPH, GSSG and GSH stable in the supernatant. The extracted sample is divided on four parts for selective measurement of the metabolites (Fig. 1a). The first aliquot is acidified to stabilize NAD^+^ and NADP^+^ while degrading their reduced counterparts. The second aliquot is alkalinized and heated to stabilize NADH and NADPH, degrading the oxidized forms. The third aliquot directly measures the combined pool of GSH and GSSG using previously published protocol with Ellman’s reagent as a reporter. The fourth aliquot measures GSSG alone, after treatment with a thiol-masking reagent to inhibit GSH detection as described before. GSH concentration is calculated by subtracting the GSSG concentration from the total glutathione pool.

### Validation of extraction conditions

The stability of NAD^+^, NADH, NADP^+^ and NADPH during the extraction was monitored using pure compounds (Fig. 1b-c) in presence and absence of blood as biological matrix. Stability of pure metabolites during extraction process in absence of biological sample was monitored by UV-VIS spectroscopy. As biological matrix can metabolize spiked pure compound we added each NAD form individually at different stages of the extraction (Supplementary Table 1) and followed recovery in enzymatic assays. When exogenous pure compounds were spiked to fresh blood (Sample Type 1), recovery rates were below 100%: NAD^+^ at 92.5%, NADH at 10%, NADP^+^ at 80% and NADPH at 60%. NADH with low recovery was detected in NAD+ assay (88%) suggesting substantial interconversion of the spiked compound. Spiking blood simultaneously with extraction buffer (Sample Type 2) resulted in improved recoveries—98% for NAD^+^, 85% for NADH (with only 10% converting to NAD^+^), 95% for NADP^+^ and 82% for NADPH. Addition of NADs post-denaturation (Sample Type 3) achieved 100% recovery for all NAD compounds. In contrast, GSH and GSSG demonstrated high stability with 95-100% recovery under all conditions.

Further characterization of metabolite compartmentalization and associated enzymatic activity revealed that blood NADs and glutathione predominantly reside within cellular fractions, with plasma containing only trace amounts of NADH and no detectable glutathione (Fig. 1d-i). Spiking experiments into the plasma fraction prior to extraction showed that enzymes catalyzing the conversion of NADH to NAD+ are localized in plasma, while those affecting NAD^+^, NADP^+^ and NADPH are associated with the cellular fraction (Supplementary Table 1, Sample Type 4).

These findings emphasize that standardization of the extraction process is crucial to minimize matrix interference, thereby ensuring reliable assay performance.

### Stability of NAD Metabolites in Blood in vitro

The stability of NAD metabolites and glutathiones was assessed in whole fresh blood samples under varying storage temperature and time. Over 24 hours at room temperature, NAD^+^ levels increased by approximately 25% (6.6µM) and NADH by 125% (0.84µM), while NADP^+^ remained stable. NADPH levels decreased to 69% of their initial concentration, a loss of 0.74 µM (Supplementary Figure 1a-f). Extending the incubation to 72 hours exacerbated these trends, and even GSH levels fell by 33%, and GSSG increased by 81% (Supplementary Figure 1g-l). These findings support limiting room temperature exposure of samples to 4 hours to prevent significant metabolite dynamics *in vitro*. Investigating the effects of freeze-thaw cycles, blood samples from 12 healthy individuals were analyzed using the NADMED method immediately post-draw and after undergoing a freeze-thaw cycle. While freezing did not impact the concentrations of NAD^+^, NADH, GSH, or GSSG, it triggered the conversion of NADPH to NADP+, with about 50±8% of NADP^+^ originating from oxidized NADPH (Supplementary Figure 1m-r). This conversion rate can be used for estimation of NADPH and NADP^+^ concentrations in fresh blood based on levels measured in frozen sample. To do so, NADP^+^ concentration measured in frozen sample can be divided on factor two. Calculated value corresponds to expected NADP^+^ level in fresh sample. To estimate NADPH concentration in the sample before freezing, calculated value is added to the NADPH concentration measured in frozen sample. A standardized thawing procedure was developed to mitigate interferences of NAD^+^ and NADH measurements in frozen samples. Samples of small volume (150-200µL) were thawed under strict cooling conditions (+0°C to +2°C) within 12-15 minutes in an ice-water bath preferentially without agitation.

### Assay performance and Validation of NADMED Assay Against Mass-Spectrometry

NADMED assays demonstrated high precision with intra-assay coefficients of variation (CV%) ranging from 2.28 to 9.02, and inter-assay precision from 3.93 to 9.37. Details on used samples number, number of aliquots and technical replicates are described in Supplementary Tables 2 and 3. The NADMED assay’s capability to detect minimal concentrations of NAD metabolites and glutathiones was quantified. Detection limits were established as >0.35 µM for NAD^+^ and NADP^+^, >0.28 µM for NADH and NADPH, >72 µM for the GSH pool and >10 µM for GSSG (Supplementary Figure 1s). These limits are well below the physiological concentrations of these metabolites in whole blood (Fig. 2), validating the NADMED assay’s effectiveness for quantitative redox profiling.

The NADMED assay was compared to targeted quantitative mass-spectrometry (MS) methods to validate its accuracy. Parallel blood aliquots from healthy controls were analyzed using the NADMED assay and two in-house MS methods: LC-MS NADomics^35^ and targeted MS-Omics^36^. Details of both MS methods are described in Targeted Metabolomics section (below). It is important to note that only the oxidized forms of NAD (NAD^+^ and NADP^+^) were included in the comparative validation due to instability of reduced forms (NADH and NADPH) in MS analyses. First, extraction step was validated by analyzing a set of parallel blood samples extracted by different methods and analyzed with MS. Both NAD^+^ and NADP^+^ were measurable from the extracts (Supplementary Figure 2u,v). Whilst NADP^+^ values were comparable between the extraction methods, NADMED extraction method resulted in significantly higher and wider spectrum of NAD^+^ values than a standard extraction method for MS. Second, detection step was validated by analyzing samples extracted with NADMED method but analyzed either by NADMED or LC-MS NADomics. Correlations between NADMED and LC-MS NADomics measurements showed good concordance for NAD^+^ and satisfactory concordance for NADP^+^ (Fig. 1j,k). This validation underscores the reliability of NADMED in matching established MS techniques, providing a robust tool for comprehensive metabolic profiling in clinical and research settings.

### Targeted Metabolomics

For validation NAD^+^ and NADP^+^ measurement in blood by NADMED method targeted LC-MS NADomics was used. This targeted NAD-specific metabolomics was developed in metabolomics facility of Finnish Institute for Molecular Medicine (FIMM) and recently published in Presterud et al^35^. For validation we used venous blood of five healthy donors. Blood was collected into 2 ml K2EDTA Vacutainers, mixed carefully by up-and-down rotation, aliquoted into 6 microtubes of 150 µL/each and frozen at −80 °C. Samples were kept frozen prior analysis. Three aliquots from each subject were analyzed by NADMED method and another three by targeted LC-MS method. For LC-MS analysis, samples were thawed on ice-water bath, then 40 µL of whole blood was injected into 450 µL cold extraction solvent comprising of mixture of Acetonitrile:Methanol:Milli-Q water in ratio 40:40:20 according to protocol described in Presterud et al^35^. Extract from each aliquot was analyzed in three technical replicates by both methods. Results were normalized on whole blood volume and compared.

Metabolomics profiling of blood extract obtained using NADMED extraction method was done by Ms-Omics laboratory, Denmark. For this experiment, we used venous K2EDTA blood collected from ten healthy subjects, e.g 10 samples. Two aliquots of 150 µL/tube were prepared for each sample and frozen at - 80°C. One aliquot of each sample was extracted using NADMED extraction method and obtained extract was frozen. Next, frozen extract and one frozen blood aliquot from each subject were sent on dry ice to Ms-Omics facility for analysis. The blood extract was analyzed as such and whole blood sample was treated with methanol to precipitate proteins followed by liquid-liquid extraction in chloroform. Organic phase used for lipidomics analysis while aqueous phase was used for semi-polar and polar metabolite analysis.

Sample analysis for semi-polar metabolites was carried out by MS-Omics as follows. The analysis was carried out using a Thermo Scientific Vanquish LC coupled to Thermo Q Exactive HF MS. An electrospray ionization interface was used as ionization source. Analysis was performed in negative and positive ionization mode. The UPLC was performed using a slightly modified version of the protocol described by Doneanu et al^36^. Peak areas were extracted using Compound Discoverer 2.0 (Thermo Scientific). Identification of compounds were performed at four levels; Level 1: identification by retention times (compared against in-house authentic standards), accurate mass (with an accepted deviation of 3 ppm), and MS/MS spectra, Level 2a: identification by retention times (compared against in-house authentic standards), accurate mass (with an accepted deviation of 3ppm). Level 2b: identification by accurate mass (with an accepted deviation of 3 ppm), and MS/MS spectra, Level 3: identification by accurate mass alone (with an accepted deviation of 3 ppm).

Sample analysis for polar metabolites was carried out by MS-Omics as follows. The analysis was carried out using a Thermo Scientific Vanquish LC coupled to Thermo Q Exactive HF MS. An electrospray ionization interface was used as ionization source. Analysis was performed in negative and positive ionization mode. The UPLC was performed using a slightly modified version of the protocol described Hsiao et al^37^. Peak areas were extracted using Compound Discoverer 2.0 (Thermo Scientific). Identification of compounds were performed at four levels; Level 1: identification by retention times (compared against in-house authentic standards), accurate mass (with an accepted deviation of 3 ppm), and MS/MS spectra, Level 2a: identification by retention times (compared against in-house authentic standards), accurate mass (with an accepted deviation of 3 ppm). Level 2b: identification by accurate mass (with an accepted deviation of 3 ppm), and MS/MS spectra, Level 3: identification by accurate mass alone (with an accepted deviation of 3 ppm).

### Handling of the samples from in-house volunteers

Blood was collected by vein puncture into 2 ml K2-EDTA Vacutainer tubes by authorized phlebomist. Aliquots of 200µL were made and placed to −80°C at time indicated in experiments. For reference range verification, the samples were frozen within 30-60 minutes from collection.

### Handling of the samples in niacin supplementation study

Study design of niacin supplementation in healthy individuals is described in detail in our previous publication^26^. During this study blood was collected every one to two weeks for follow-up measurements. Blood was collected into 6 ml K2-Vacutainer and separated on Ficoll 400 Density gradient according to manufacturer instructions. Volumes of recovered erythrocytes fraction, containing target metabolites, varied within 40%-45% of initial blood sample resulting in x2,5-2,2 concentration factors, respectively. Plasma and erythrocytes fraction were collected and provisionally placed at −80°C until analysis. Redox profiling was done for both plasma and red blood cells using NADMED methodology. Results for red blood cell were normalized per protein content in the samples, e.g nmol/mg of protein. Protein concentration was measured using BCA kit (Pierce) according to manufacturer’s instructions. As for every sample volume of erythrocyte fraction was known, it gave us possibility to calculate concentration of NADs and glutathione in initial blood sample before Density gradient separation. To do so, measured µM concentration of target metabolite in the erythrocyte fraction was divided on concentration factor after gradient.

For muscle NAD^+^ and other correlation analyses, the data collected during our previous study^27^ was revisited.

### Sample Handling in Red Cross Blood Service

Whole blood samples of healthy donors were obtained from the Blood Service Biobank in Finland. Donors were informed about the study and gave a written consent prior blood donation for the study. Altogether 300 samples were collected, but the consent for one sample was retracted resulted in 299 reference samples.

Blood donation in Finland is tightly regulated regarding health and lifestyle for the donors. The list of restrictions is published on website www.bloodservice.fi. Age of donors varied between 18 and 70 years. Weight of a donor was within 50-200 kg. Blood for this study was collected during days 9.12.2019, 27.01.2020, 4.02.2020, 5.02.2020, 10.02-11.02.2021.

Venous blood was first collected into a pouch of a blood donation unit containing trace amounts of citrate. Sample for the study was taken from the pouch into 4 ml K2-EDTA tube. Samples were transported to central BioBank facility at ambient temperature in the time interval from 3 to 17 hours. Then 200 ul aliquot was taken into a small tube, sealed and kept at +4 C° for additional time interval of 8 to 15 hours followed by transferring into −20 C° freezer. In summary, time interval between blood withdrawal and freezing varied from 11 to 32 hours. After keeping samples for 48h at −20 C° they were transferred into −80 C° freezer. Samples were delivered on dry ice for analysis. All samples obtained by the Blood Service Biobank were in frozen 200 ul aliquots, anonymized, coded and accompanied with gender, age, height and weight information.

### Sample Handling in Turku University Hospital Auria BioBank Pathology Collection

All blood samples were taken from BioBank storage after given informed consent from patients. Venous whole blood samples from patients were collected into 10 ml K2EDTA Vacutainers, centrifuged at 1000g for 10 min at +4°C to collect 1,8 ml of plasma. After removal of target volume of plasma, samples were carefully mixed again resulting in x1,22 concentrated blood. Next, 400 µl of concentrated blood was pipetted into a small tube, sealed and frozen at −80°C.

Time interval between sample withdrawal and freezing was within 1,5-8 hours with average of 5 hours. During this time, samples were kept at room temperature. Ten samples were different from the rest as they were kept for 25-28 hours before processing and freezing. Auria BioBank samples were coded and accompanied with information about gender, age, diagnosis and time interval between diagnosis date and sample collection.

### Chemicals

All reagents used in the study were from Sigma and of highest purity available.

### Data analysis

Data analysis and its graphical representation was done using GraphPad Prism version 9 and 10. Statistical analyses used are specified in the legend of each figure. Outliers were removed from control data after identification with ROUT Q = 0.5% (number of outliers removed: NAD^+^/NADH ratio n = 2, NADP^+^/NADPH ratio n = 11 and GSH/SSG ratio n = 6, no outliers were identified for individual metabolites).

### Patent portfolio

Euro, L. and Wartiovaara, A. (2023) ‘METHOD FOR DETERMINING AMOUNTS OF NAD METABOLITES FROM SAMPLE AND METHODS AND USES RELATED THERETO’. EP: HELSINGIN YLIOPISTO OP - FI 20205738 A OP - FI 2021050529 W. Available at: https://lens.org/055-672-380-749-328.

Euro, L. and Wartiovaara, A. (2022) ‘METHOD FOR DETERMINING AMOUNTS OF NAD METABOLITES FROM SAMPLE AND METHODS AND USES RELATED THERETO’. WO: HELSINGIN YLIOPISTO OP - FI 20205738 A. Available at: https://lens.org/154-826-164-167-292.

Euro, L. and Wartiovaara, A. (2023a) ‘Method for determining amounts of NAD metabolites from sample and methods and uses related thereto’. AU: HELSINGIN YLIOPISTO OP - FI 20205738 A OP - FI 2021050529 W. Available at: https://lens.org/023-253-113-887-235.

Euro, L. and Wartiovaara, A. (2023b) ‘METHOD FOR DETERMINING AMOUNTS OF NAD METABOLITES FROM SAMPLE AND METHODS AND USES RELATED THERETO’. US: HELSINGIN YLIOPISTO OP - FI 20205738 A OP - FI 2021050529 W. Available at: https://lens.org/166-870-053-588-047.

