## Supplementary Material for "Dynamics of blood NAD and glutathione in health, disease, aging and under NAD-booster treatment"

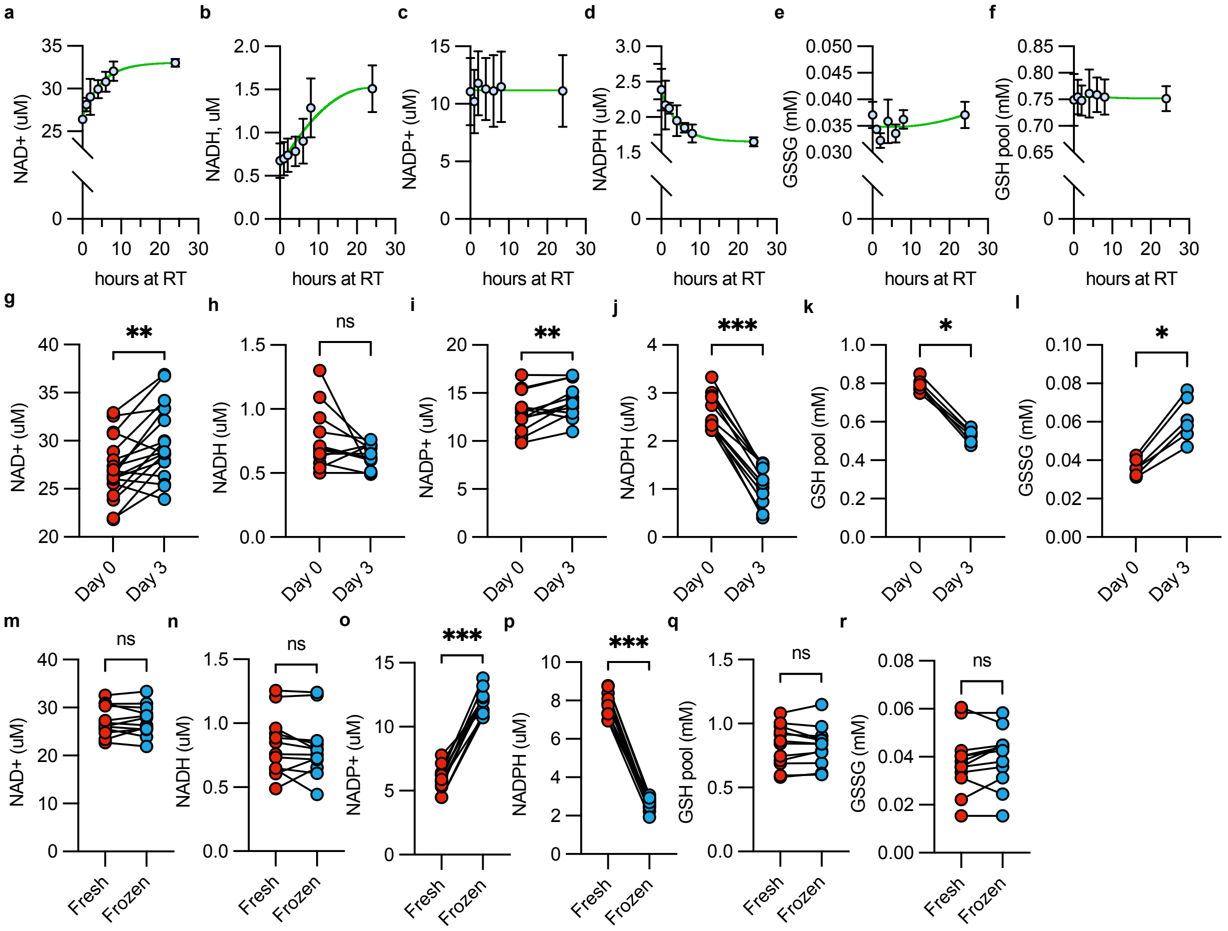

**s**

Practical Quantitation Limit

| Metabolite | uM in whole blood |
|---|---|
| NAD(P)+ | 0.35 |
| NAD(P)H | 0.28 |
| GSH pool | 72 |
| GSSG | 10 |

**Supplementary Figure 1. Properties of blood NAD metabolites and glutathiones in fresh and frozen blood.**

**a-f.** Stability of NAD+ (a), NADH (b), NADP+ (c), NADPH (d), GSH pool (e) and GSSG (f) (n=4, 4, 5, 5, 3 and 2 per timepoint, respectively) in whole blood during 24 h storage after withdrawal at ambient temperature before freezing. **g-l.** Stability of NAD+ (n=18) (g), NADH (n=12) (h), NADP+ (n=12) (i), NADPH (n=12) (j), GSH pool (n=6) (k) and GSSG (n=6) (l) in whole blood during three-day storage after withdrawal at ambient temperature before freezing. **m-r.** Levels of NAD+ (m), NADH (n), NADP+ (o), NADPH (p), GSH pool (q) and GSSG (r) before and after freezing the sample (n=12). Metabolite extraction from one aliquot of the sample and freezing of the other aliquot was done at the same time after blood withdrawal, thus within 30 min. **s.** Practical quantitation limits for each assay.

Wilcoxon matched-pairs signed rank test used when p-values given. *$p < 0.05$; **$p < 0.01$; ***$p < 0.001$

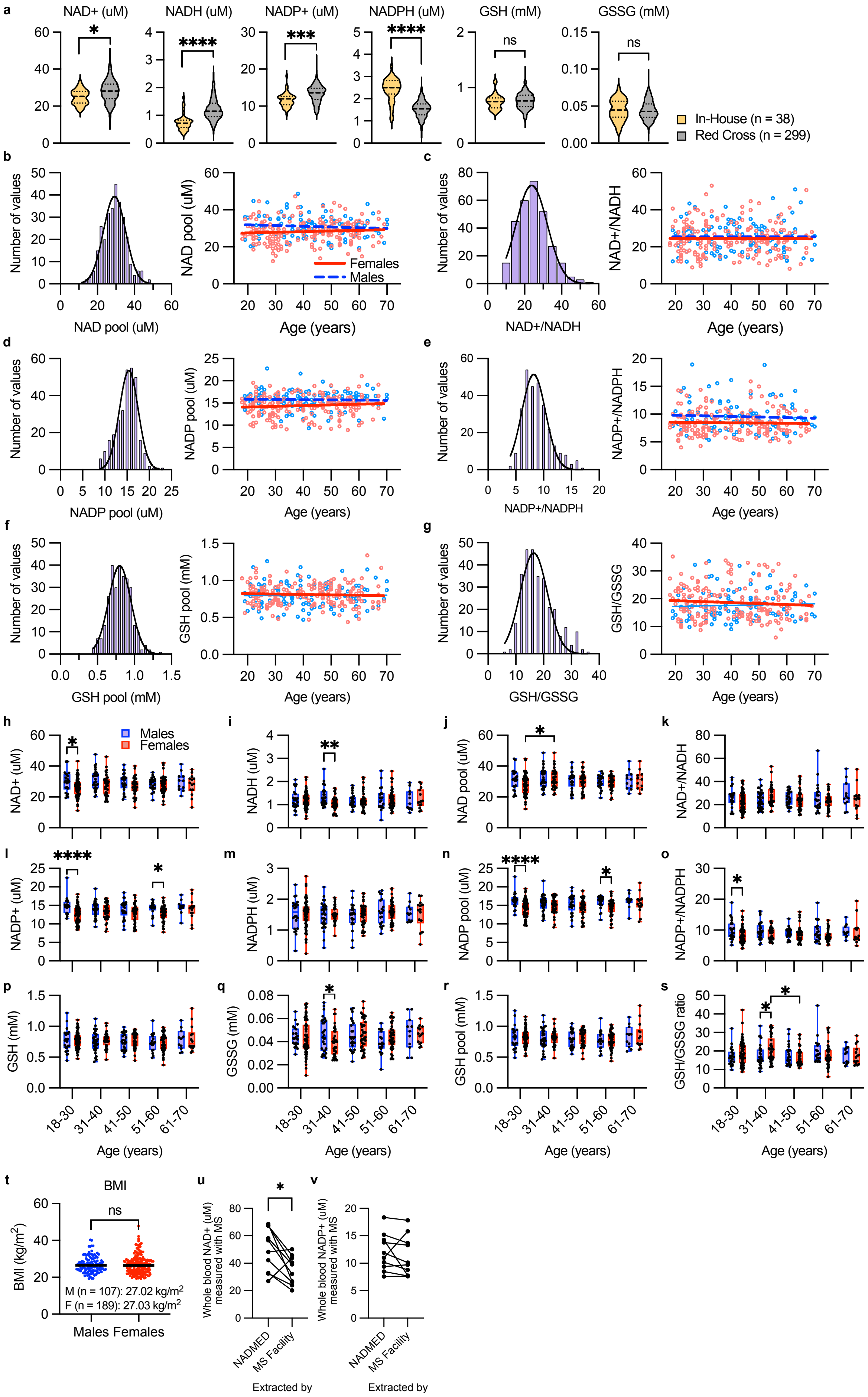

a
NAD+ (uM)
NADH (uM)
NADP+ (uM)
NADPH (uM)
GSH (mM)
GSSG (mM)
*
****
***
****
ns
ns
In-House (n = 38)
Red Cross (n = 299)
b
Number of values
NAD pool (uM)
Females
Males
Age (years)
c
NAD+/NADH
d
NADP pool (uM)
e
NADP+/NADPH
f
GSH pool (mM)
g
GSH/GSSG
h
NAD+ (uM)
Males
Females
i
NADH (uM)
j
NAD pool (uM)
k
NAD+/NADH
l
NADP+ (uM)
m
NADPH (uM)
n
NADP pool (uM)
o
NADP+/NADPH
p
GSH (mM)
q
GSSG (mM)
r
GSH pool (mM)
s
GSH/GSSG ratio
18-30
31-40
41-50
51-60
61-70
Age (years)
t
BMI
ns
BMI (kg/m2)
M (n = 107): 27.02 kg/m2
F (n = 189): 27.03 kg/m2
Males Females
u
Whole blood NAD+ (uM) measured with MS
NADMED
MS Facility
Extracted by
v
Whole blood NADP+ (uM) measured with MS
NADMED
MS Facility
Extracted by

**Supplementary Figure 2. Pools and ratios of NAD metabolites and glutathiones in blood of healthy population remain stable during aging and follow normal distribution.**

**a.** Whole blood NAD metabolite and glutathione levels in in-house controls and Red Cross blood donors. Populations differ on their sampling-to-freezing time with in-house control samples frozen within an hour while Red Cross samples were frozen within approximately 10-30 hours. **b-g.** Frequency distribution and levels of NAD pool (b), NAD+/NADH ratio (c), NADP pool (d), NADP+/NADPH ratio (e), GSH pool (f) and GSH/GSSG ratio (g) in the whole blood of 299 healthy donors at the age of 18-70 years. Red and blue lines represent linear regression fit of the data separately for females and males. Black line represents Gaussian distribution. **h-s.** Levels of NAD metabolites and glutathione and their pools and rations in the whole blood in males and females separated to different age groups. **t.** BMIs of males and females. BMI was not available from two individuals, and they are excluded from the analysis. **u and v.** Comparison of NAD+ (u) and NADP+ (v) in whole blood samples with different extraction methods. Metabolite levels measured by MS Facility.

In panel a, Mann-Whitney nonparametric t-test was applied. In panels b-t, metabolite levels were measured from 299 healthy blood donors at age of 18-70 years. During extraction of GSH and GSSG one sample was lost. Two-way ANOVA with Tukey's multiple comparison test was used in panels h-s. Unpaired t-test was used in graph t. Wilcoxon matched-pairs signed rank test used in graphs u and v.

| Spike with§ | | NAD+ | | NADH | | NADP+ | | NADPH | | GSH | GSSG |
|---|---|---|---|---|---|---|---|---|---|---|---|
| Average recovery (%) as | | NAD+ | NADH | NAD+ | NADH | NADP+ | NADPH | NADP+ | NADPH | GSH | GSSG |
| Sample type | 1. Fresh/thawed blood before extraction | 92.5 | ND | 88 | 10 | 80 | ND | ND | 60 | 95 | 98 |
| | 2. Fresh/thawed blood during hot extraction | 98 | ND | 10 | 85 | 95 | ND | ND | 82 | 97 | 98 |
| | 3. Homogenate | 100 | ND | ND | 100 | 100 | ND | ND | 100 | | |
| | 4. Plasma before extraction | 100 | ND | 98 | ND | 100 | ND | ND | 95 | 97 | 100 |

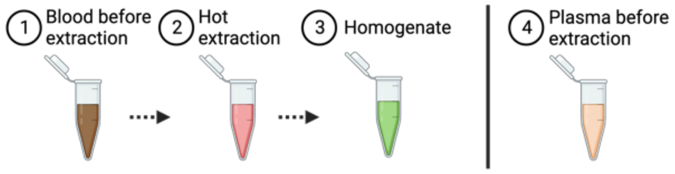

**Supplementary Table 1. Recovery of pure components spiked to the sample at different stages during extraction.** Spiked amounts were in range of physiological concentrations of NAD metabolites in blood – 1,4, 10, 25 uM. Spiking experiments were done at different times and by two operators.

Intra-Assay precision

| | NAD+ | | | NADH | | | NADP+ | | | NADPH | | | GSH pool | | | GSSG | | |
|---|---|---|---|---|---|---|---|---|---|---|---|---|---|---|---|---|---|---|
| Sample ID | 1 | 2 | 3 | 1 | 2 | 3 | 1 | 2 | 3 | 1 | 2 | 3 | 1 | 2 | 3 | 1 | 2 | 3 |
| Mean (uM/mM) | 27.41 | 29.41 | 22.00 | 0.55 | 0.71 | 0.64 | 10.03 | 11.59 | 11.01 | 2.39 | 2.57 | 2.62 | 1.03 | 0.86 | 0.76 | 0.03 | 0.022 | 0.036 |
| SD (uM/mM) | 0.62 | 1.31 | 0.87 | 0.03 | 0.05 | 0.05 | 0.83 | 0.58 | 0.54 | 0.16 | 0.09 | 0.10 | 0.05 | 0.04 | 0.03 | 0.002 | 0.002 | 0.003 |
| CV (%) | 2.28 | 4.45 | 3.95 | 5.20 | 7.06 | 8.45 | 8.28 | 4.98 | 4.90 | 6.50 | 3.67 | 3.81 | 5.09 | 4.27 | 3.88 | 5.82 | 6.97 | 9.02 |

**Supplementary Table 2. Intra-assay Precision (precision within an assay) for NAD+, NADH, NADP+, NADPH, GSH pool and GSSG.** Whole blood from three healthy subjects was used (sample ID 1-3). Three aliquots of each sample were extracted and obtained stabilized extracts were analyzed in three or four technical replicates for each metabolite.

Inter-Assay precision

| | NAD+ | NADH | NADP+ | NADPH | GSH pool | GSSG |
|---|---|---|---|---|---|---|
| Mean (uM/mM) | 29.36 | 0.76 | 10.91 | 2.35 | 0.782 | 0.031 |
| SD (uM/mM) | 1.15 | 0.07 | 0.94 | 0.14 | 0.056 | 0.003 |
| CV (%) | 3.93 | 9.37 | 8.59 | 5.91 | 7.18 | 8.51 |

**Supplementary Table 3. Inter-assay Precision (precision between assays).** For Inter-Assay Precision blood sample from one healthy subject was used. Sample was aliquoted and frozen. Individual aliquots were measured in six separate assays for each metabolite using 10 technical replicates in every assay.

| | | 18-30 years | | | | 31-40 years | | | | 41-50 years | | | | 51-60 years | | | | 61-70 years | | | | RedCross donors | | In-house controls (frozen within 30 min) | |
|---|---|---|---|---|---|---|---|---|---|---|---|---|---|---|---|---|---|---|---|---|---|---|---|---|---|
| | | Males | | Females | | Males | | Females | | Males | | Females | | Males | | Females | | Males | | Females | | 18-70 years | | 25-65 years | |
| N | | 22 | | 64 | | 30 | | 33 | | 25 | | 39 | | 20 | | 41 | | 11 | | 14 | | 298-299 | | 32-38 | |
| Metabolite | | Mean | ± SD | Mean | ± SD | Mean | ± SD | Mean | ± SD | Mean | ± SD | Mean | ± SD | Mean | ± SD | Mean | ± SD | Mean | ± SD | Mean | ± SD | Mean | ± SD | Mean | ± SD |
| uM | NAD+ | 29.82 | 6.64 | 26.09 | 5.94 | 30.90 | 6.56 | 27.93 | 6.84 | 29.42 | 5.91 | 26.88 | 5.83 | 28.48 | 4.40 | 27.75 | 5.72 | 29.95 | 6.16 | 27.89 | 6.76 | 28.05 | 6.18 | 24.87 | 3.96 |
| | NADH | 1.22 | 0.41 | 1.21 | 0.36 | 1.35 | 0.39 | 1.05 | 0.31 | 1.20 | 0.30 | 1.18 | 0.37 | 1.27 | 0.49 | 1.26 | 0.41 | 1.20 | 0.45 | 1.25 | 0.42 | 1.22 | 0.39 | 0.7514 | 0.288 |
| | NAD pool | 31.05 | 6.71 | 27.30 | 5.96 | 32.25 | 6.62 | 29.02 | 7.04 | 30.62 | 6.03 | 28.06 | 5.85 | 29.75 | 4.51 | 29.01 | 5.89 | 31.15 | 6.20 | 29.14 | 6.73 | 29.26 | 6.27 | 25.62 | 4.078 |
| | NAD+/NADH | 26.50 | 8.52 | 23.22 | 8.26 | 24.30 | 7.61 | 27.69 | 8.79 | 25.45 | 6.23 | 24.79 | 8.88 | 24.13 | 8.91 | 23.50 | 6.44 | 28.36 | 11.91 | 25.02 | 11.01 | 24.83 | 8.35 | 37.07 | 13.86 |
| uM | NADP+ | 14.85 | 2.25 | 12.51 | 2.27 | 14.02 | 2.17 | 13.27 | 2.31 | 13.88 | 2.47 | 12.91 | 2.53 | 14.17 | 1.77 | 12.80 | 2.08 | 14.61 | 1.92 | 14.03 | 2.78 | 13.38 | 2.37 | 11.74 | 1.936 |
| | NADPH | 1.52 | 0.50 | 1.53 | 0.45 | 1.49 | 0.42 | 1.51 | 0.26 | 1.47 | 0.40 | 1.50 | 0.35 | 1.61 | 0.43 | 1.59 | 0.33 | 1.57 | 0.30 | 1.53 | 0.47 | 1.53 | 0.39 | 2.408 | 0.5297 |
| | NADP pool | 16.36 | 2.10 | 14.04 | 2.37 | 15.51 | 2.27 | 14.78 | 2.28 | 15.34 | 2.53 | 14.42 | 2.58 | 15.78 | 1.78 | 14.40 | 2.19 | 16.18 | 1.97 | 15.56 | 2.71 | 14.91 | 2.41 | 14.15 | 2.007 |
| | NADP+/NADPH | 10.01 | 3.28 | 8.46 | 2.65 | 9.70 | 2.77 | 8.78 | 2.03 | 9.03 | 1.75 | 8.67 | 2.53 | 9.54 | 3.32 | 8.31 | 2.00 | 9.56 | 2.11 | 9.43 | 4.17 | 8.87 | 2.47 | 4.786 | 0.9396 |
| mM | GSH | 0.77 | 0.19 | 0.77 | 0.13 | 0.75 | 0.17 | 0.77 | 0.12 | 0.75 | 0.13 | 0.77 | 0.12 | 0.73 | 0.14 | 0.73 | 0.15 | 0.78 | 0.17 | 0.80 | 0.21 | 0.76 | 0.15 | 0.7484 | 0.1347 |
| | GSSG | 0.04 | 0.01 | 0.04 | 0.01 | 0.05 | 0.01 | 0.04 | 0.01 | 0.04 | 0.01 | 0.05 | 0.01 | 0.04 | 0.01 | 0.04 | 0.01 | 0.05 | 0.01 | 0.05 | 0.01 | 0.04 | 0.01 | 0.04 | 0.01 |
| | GSH pool | 0.82 | 0.19 | 0.82 | 0.13 | 0.79 | 0.17 | 0.81 | 0.12 | 0.79 | 0.14 | 0.82 | 0.13 | 0.77 | 0.15 | 0.78 | 0.15 | 0.83 | 0.18 | 0.85 | 0.21 | 0.81 | 0.15 | 0.79 | 0.14 |
| | GSH/GSSG | 17.18 | 4.90 | 18.88 | 6.42 | 17.46 | 5.87 | 21.16 | 6.73 | 17.72 | 5.43 | 16.99 | 4.72 | 18.51 | 4.12 | 17.73 | 6.10 | 17.63 | 4.51 | 17.78 | 5.16 | 18.26 | 5.71 | 17.35 | 5.251 |

**Supplementary Table 4. Calculated mean values ± standard deviation for all measured metabolites in blood samples of 299 Red Cross blood donors and 38 in-house controls.** For GSH and GSSG 298 samples from Red Cross donors and 32 samples from in-house controls could be analyzed.

| | | Estimated reference range for fresh blood | In-house controls (fresh) |
|---|---|---|---|
| | N | 337 | 12 |
| uM | NAD+ | 21.7 - 33.7 | 24.1 - 30.44 |
| uM | NADH | 0.8 - 1.6 | 0.59 - 1.06 |
| uM | NAD pool | 22.7 - 35.1 | 23.53 - 32.67 |
| | NAD+/NADH | 16.6 - 35.8 | 25.03 - 45.03 |
| uM | NADP+ | 4.9 - 7.9 | **4.91 -6.91** |
| uM | NADPH | 6.6 - 10 | **7.08 - 8.38** |
| uM | NADP pool | 12.1 - 17.3 | **12.24 - 15.04** |
| | NADPH/NADP | 1.19 - 1.42 | **1.12-1.54** |
| mM | GSH | 0.62 - 0.91 | 0.64 - 0.94 |
| mM | GSSG | 0.03 - 0.05 | 0.03 - 0.05 |
| mM | GSH pool | 0.65 - 0.95 | 0.67 - 0.99 |
| | GSH/GSSG | 12.5 - 23.9 | 16.4 - 28.4 |

**Supplementary Table 5. Estimated control ranges for each redox metabolite in fresh blood.** Right most panel represents values from 12 in-house controls whose blood samples were analyzed fresh. Intervals calculated as mean +/- standard deviation.

| Diagnosis | ICD-10 | n | Days from diagnosis to sampling | | | | | | | |
|---|---|---|---|---|---|---|---|---|---|---|
| | | | **-450...-250** | **-249...-50** | **-49...-1** | **+1...+35** | **+36...+100** | **+101...+250** | **+251...+450** | **+451...+4050** |
| Malignant neoplasm of pancreas | C25 | 15 | | 2 | 2 | 8 | | 3 | | |
| Malignant neoplasm of breast | C50 | 22 | | 7 | 15 | | | | | |
| Acute myelogenous leukemia (AML) | C92.0 | 12 | | | | | | 2 | 1 | 9 |
| Diabetes type II | E11 | 15 | | 5 | 10 | | | | | |
| Parkinson disease | G20 | 15 | | 5 | | 1 | 4 | 4 | 1 | |
| Alzheimer disease | G30 | 15 | | 4 | 4 | 1 | 4 | 2 | | |
| Prostate adenocarcinoma | C61 | 14 | | 5 | 9 | | | | | |
| Skin malignancy | C44 | 15 | | 8 | 7 | | | | | |
| Ovarian epithelial cancer | C56 | 12 | 2 | 2 | 5 | 1 | 1 | 1 | | |
| Malignant neoplasm of colon | C18 | 14 | | 6 | 8 | | | | | |
| Malignant neoplasm: upper lobe, bronchus or lung tumor | C34.1 | 15 | | 4 | 11 | | | | | |
| | Total | 164 | 121 | | | 43 | | | | |

**Supplementary Table 6. Time of blood sample collection in relation to disease diagnosis.** Disease groups with only pre-diagnosis samples highlighted with light gray.
